# Relative-location tuning in five frontal areas in a rhesus monkey

**DOI:** 10.1101/2025.10.25.684521

**Authors:** Mikhail A. Lebedev, Steven P. Wise

**Affiliations:** Faculty of Mechanics and Mathematics, Lomonosov Moscow State University, Moscow, Russia; Research Center in the Field of Artificial Intelligence, Lomonosov Moscow State University, Moscow, Russia; Sechenov Institute of Evolutionary Physiology and Biochemistry of the Russian Academy of Sciences, St. Petersburg, Russia; Olschefskie Institute for the Neurobiology of Knowledge, Bethesda, MD 20814, USA

## Abstract

Primates exhibit remarkable flexibility in coordinating gaze angle, spatial attention, and motor goals, such as fixating on one point, attending to another, and planning a reach toward a third. Neuronal reference frames are critical for these behaviors. We investigated relative-location tuning in macaque monkeys, where the reference frame was defined by the relative positions of two objects in a scene rather than their absolute coordinates. Single neurons were recorded from five frontal cortex regions: dorsal and ventral premotor cortex (PMd and PMv), supplementary motor area (SMA), dorsolateral prefrontal cortex (PFdl), and frontal eye field (FEF). In each trial, the monkey viewed two food pellets, one in a stationary feeder and another in a feeder attached to a moving robot, which served as an attention attractor. By task rule, the monkey could obtain only one pellet. A significant proportion of neurons in all five regions showed spatial tuning based on the relative positions of the two feeders (∼35-65%, depending on the area). For instance, some neurons were more active when one feeder was positioned to the right of the other. In PFdl and FEF, most relative-location neurons primarily encoded the monkey’s fixation on the relatively left or right feeder. In SMA, PMd, and PMv, some neurons reflected fixation, but more encoded whether the relatively left or right feeder was the attention attractor (independent of gaze) or the reach target. Relative-location tuning was more prevalent when both feeders were task-relevant, such as when the monkey attended to one feeder while planning a reach to the other. Relative-location neurons could contribute to computing object-based reference frames or allocating spatial information-processing resources across attentional, oculomotor, and skeletomotor behaviors. They also suffice to explain gaze-dependent reach-related activity without contradicting theories of visually guided reaching, which predict gaze invariance.

## Introduction

Primates, including humans, demonstrate flexibility in spatially guided behavior. Typically, they fixate on the target of a reaching movement while allocating significant attentional resources to it, aligning the reach target, spatial attention, and gaze fixation. However, primates can also prepare a reach toward one location, attend to another, and fixate on a third. In our 2001 study, we reported that different types of neurons in the dorsal premotor cortex (PMd) are preferentially attuned to specific spatial parameters such as planned and executed movements, attention, or gaze (Lebedev and Wise, 2001). Here, we extend the analysis of the same data to investigate relative-location coding of these spatial variables in PMd and four additional frontal cortex regions.

Relative-location neuronal coding has been documented in previous studies. An early study in behavioral neurophysiology found that neurons in the dorsolateral prefrontal cortex (PFdl) of monkeys encode the relative positions of stimuli rather than their absolute locations in visual space (Niki, 1974). More recent research has shown that neurons in the supplementary eye field (SEF), another frontal cortex region, encode the position of one part of an object relative to another (Olson and Gettner, 1995; Olson and Tremblay, 2000; Tremblay et al., 2002). Relational coding could represent a general property of the frontal cortex, extending beyond spatial domains. For instance, neurons in the orbital prefrontal cortex encode the relative value of food items (Tremblay and Schultz, 1999; Hikosaka and Watanabe, 2000).

Neurons in the posterior parietal cortex (PPC) also exhibit relative coding of spatial locations, though this is typically interpreted within a different theoretical framework. Andersen and colleagues demonstrated that some PPC neurons involved in reaching encoded the target’s location relative to both the monkey’s hand and eye (Buneo et al., 2002). They proposed that the PPC uses a retinocentric reference frame during the planning and execution of reaching movements. Furthermore, the high-level plan for a reach corresponds to a difference vector in retinocentric space, representing the displacement between the current hand position and the target location.

The other groups reported that neuronal activity in the ventral premotor cortex (PMv) remained invariant to eye position (Fogassi et al., 1992, 1996; Graziano and Gross, 1998) and hand orientation (Kakei et al., 2001; see also Hoshi and Tanji, 2002) but varied with hand location (Graziano et al., 1997). Many of these early studies relied on semi-quantitative, receptive-field-like analyses of neuronal activity. In contrast, quantitative analyses consistently revealed significant effects of gaze orientation on neuronal activity in PMv (Boussaoud et al., 1993; Mushiake et al., 1997), dorsal premotor cortex (PMd) (Boussaoud et al., 1993; Boussaoud, 1998; Boussaoud and Bremmer, 1999; Cisek and Kalaska, 2002), supplementary motor area (SMA), and pre-supplementary motor area (Fujii et al., 2002). Similar gaze-dependent effects were observed in the posterior parietal cortex (PPC) (Andersen et al., 1990; Brotchie et al., 1995; Battaglia-Mayer et al., 2000, 2001; Ferraina et al., 2001).

Experiments demonstrating gaze dependency of reach-related signals in PMv and PMd all required monkeys to fixate on one location while planning and executing a reach, typically to a distinct target. In this study, we expanded this paradigm to investigate whether spatial attention, independent of gaze direction, modulates neuronal spatial tuning. Specifically, we examined neurons encoding the relative positions of task-relevant objects in the scene and analyzed how their tuning depends on gaze direction, spatial attention, and reach preparation.

## Materials and Methods

The data reported here was collected in 1998-2000. One male rhesus monkey (*Macaca mulatta*), weighing 7.0 kg, was studied in accord with the *Guide for the Care and Use of Laboratory Animals* (1996). The animal care and experimental procedures were approved by the National Institute of Mental Health Animal Care and Use Committee. The same monkey was the subject of the report, which dealt with a different aspect of neuronal activity in PMd (Lebedev and Wise, 2001).

We manipulated spatial attention, gaze orientation, and reach target using an experimental design featuring a robotically controlled food dispenser. The monkey attended to the robot’s movements to determine when to initiate a reach. The reach target was either a food pellet moved by the robot (Task 1) or a stationary pellet in a fixed feeder at a different location (Task 2).

### Experimental apparatus and behavioral tasks

During the experiment, the monkey sat in a primate chair with its head fixed and left arm restrained. A touch pad was attached to the chair at waist level on the monkey’s right side. An oculometer (Bouis Instruments, Karlsruhe, Germany) was placed in front of the right eye, and eye position was sampled at 250 Hz. Two food-pellet dispensers (Mitz et al., 2001), or feeders, were located in front of the monkey (Fig. 1). One of them, termed the robot feeder (R), was positioned by a robot (Sands Technology, Cambridge, UK). The other feeder was manually positioned, and thus was termed the manual feeder (M). The R’s pellet started beyond the reach of the monkey, and only when it first moved within reach could the monkey retrieve one of the pellets. The monkey, therefore, had to allocate considerable attentional resources to the location of the R. The monkey performed two tasks. In one, the monkey could only get the R’s pellet; in the other task, only the M’s pellet was available. The rule for pellet availability (R vs. M) varied in blocks of trials.

**Figure 1.**
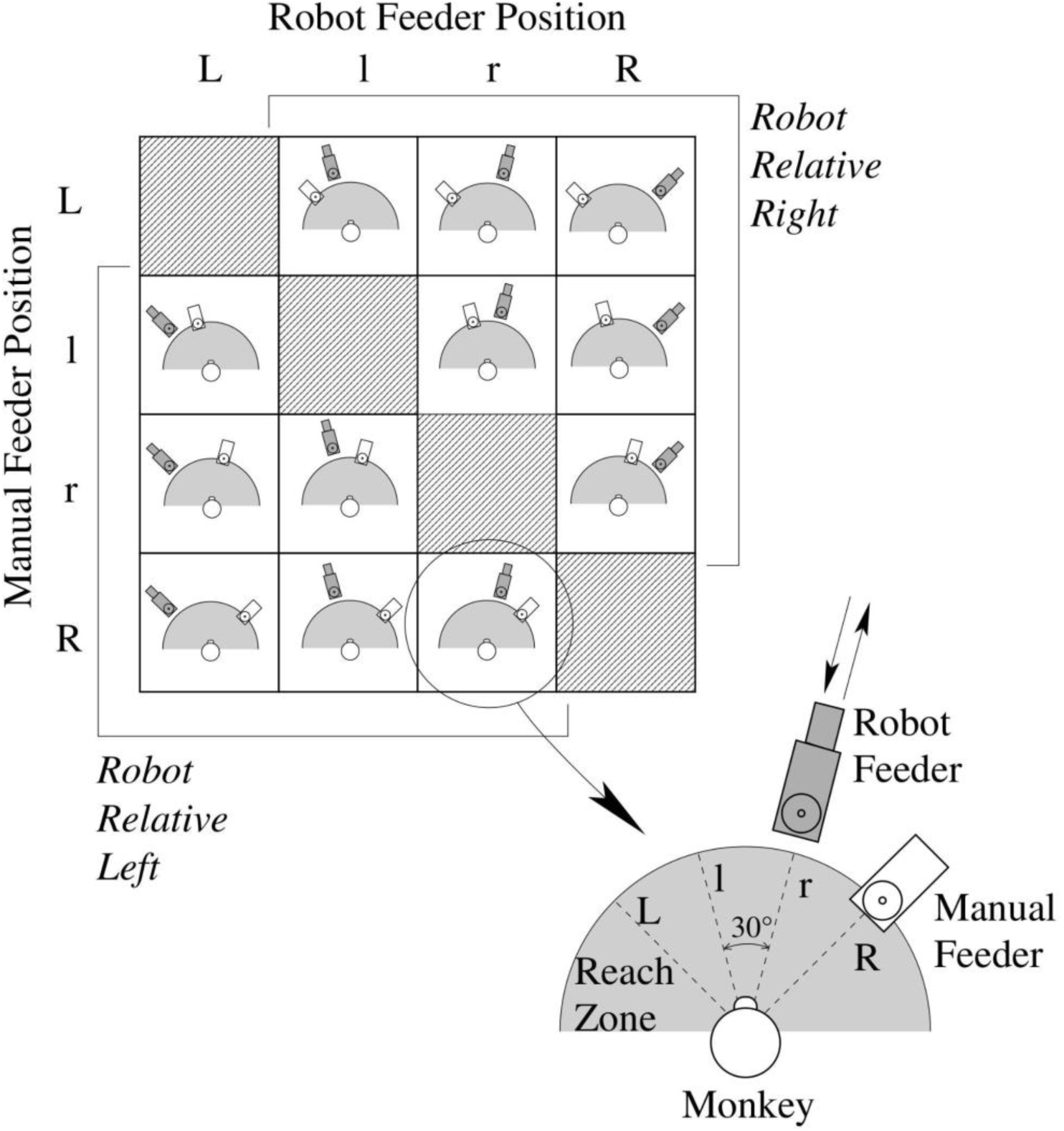
Feeder locations represented as a matrix. Possible arrangements of the feeders around the monkey are shown as views from above. One arrangement is shown in more detail at the bottom right. Four positions were designated for the robot and manual feeders, marked L (far left), l (near left), r (near right) and R (far right). The manual feeder remained inside the zone reachable by the monkey (shaded). The robot feeder started outside of the monkey’s reach and, after a random sequence of movements back and forth, entered the reach zone to signal that the monkey could reach for a food pellet. The major diagonal of the matrix (hatched) did not correspond to any valid configuration of the feeders because the two feeders could not be physically placed at the same location. Matrix elements above and below the major diagonal correspond to the robot on the right and on the left from the manual feeder, respectively (robot relative right versus robot relative left).

In both tasks, the monkey placed its right hand on the touch pad to begin a trial. The monkey had to maintain contact with the touch pad until much later in the trial, when a movement-triggering event occurred. Once the monkey’s hand contacted the touch pad, the food pellets (45-mg cereal pellet, Primate Products, Woodside, California, USA) were delivered automatically to each feeder. The M’s pellet was always within reach of the monkey, but the R’s pellet came into reach only once per trial to signal that the monkey could reach toward a pellet (although not necessarily the pellet moved by the robot). The monkey could obtain only one food pellet on each trial. By rule imposed by the experimental design, in Task 1, the monkey could retrieve only the R’s pellet; in Task 2, it could get only the M’s pellet. If the monkey reached for the disallowed pellet, its hand’s approach to the pellet dispenser was sensed by a proximity detector, which opened a trap door supporting the food pellet, and the pellet fell away before the monkey could acquire it. If the monkey attempted to reach at a disallowed time, the pellets were removed via the same trap-door mechanism immediately after the monkey lifted its hand from the touch pad.

On each trial, the R was set in motion 2.4 s after trial onset, moving alternatively toward and away from the monkey. Each motion had a randomly selected amplitude of 4.5–9.0 cm and a duration of 0.37–0.60 s. Every R movement was separated by 0.5–1.4 s pauses (unpredictably varied), and the number of robot movements ranged from 1 to 6 on any given trial. During robot movements, the R’s pellet remained beyond the monkey’s reach (by up to 9 cm), except for the final movement, which was indistinguishable from the others in amplitude and duration. This final movement started from a random location at an unpredictable time, and it brought the R’s pellet a distance of 4.5 cm within a perimeter reachable by the monkey, termed the *reach zone* (Fig. 1). When the R’s pellet entered the reach zone, the monkey needed to reach the appropriate (*i.e.,* allowed) feeder within 300 ms. If the monkey reached the correct feeder within that time, its pellet remained and the other pellet fell away. The R stopped moving as the monkey retrieved the pellet.

Task 1 (“reach to the R”) and Task 2 (“reach to the M”) alternated in blocks of 12–24 trials. The R and M were restricted, by design, to four locations. Fig. 1 illustrates those locations, separated by 30° and termed far left (abbreviated L), near left (l), near right (r), and far right (R). This range of locations was selected because they could be easily fixated by the head-fixed monkey. In each block of trials, the M remained in one of the four positions. Task 1 began first. The robot changed the R’s location every 4–8 trials until all of three unoccupied locations had been used. The R then returned to its initial location for the block, which signaled that Task 2 had begun, and the same sequence of R locations was repeated. Note that the return of the R to its initial location for a block provided the monkey with a highly conspicuous visual cue about when to change from Task 1 to Task 2. This second sequence of R movements concluded a block of trials, after which the M was repositioned, and Task 1 resumed. Note that moving the M provided the monkey with a clear cue about when to change from Task 2 to Task 1. Recording of a neuron’s activity continued until all four M positions had been used 1–3 times each.

### Analysis of neuronal activity

The discharges of individual neurons were recorded using either a conventional single-electrode microdrive or a 7-electrode drive (Uwe Thomas Recordings, Marburg, Germany). Electrode impedance was 0.5-1.5 MΩ, measured at 1 KHz. After amplifying and filtering with a band pass of 0.6 to 6.0 KHz, single-unit potentials were discriminated using the Multispike Detector (Alpha-Omega Engineering, Nazareth, Israel) or a time-amplitude waveform discriminator (BAK Electronics, Rockville, Maryland). The number of individual neurons isolated from SMA, PMd, PMv, FEF, and PFd was 135, 199, 67, 19 and 83, respectively.

The monkey’s oculomotor behavior was not instrumentally conditioned. However, the monkey typically fixated one of the two most salient features in its environment: the R or the M (see eye-position records in Fig. 2A, B). Stable fixation epochs of at least 0.3 s were selected for analysis, with the initial and final 0.1 s discarded for the final measure.

**Figure 2.**
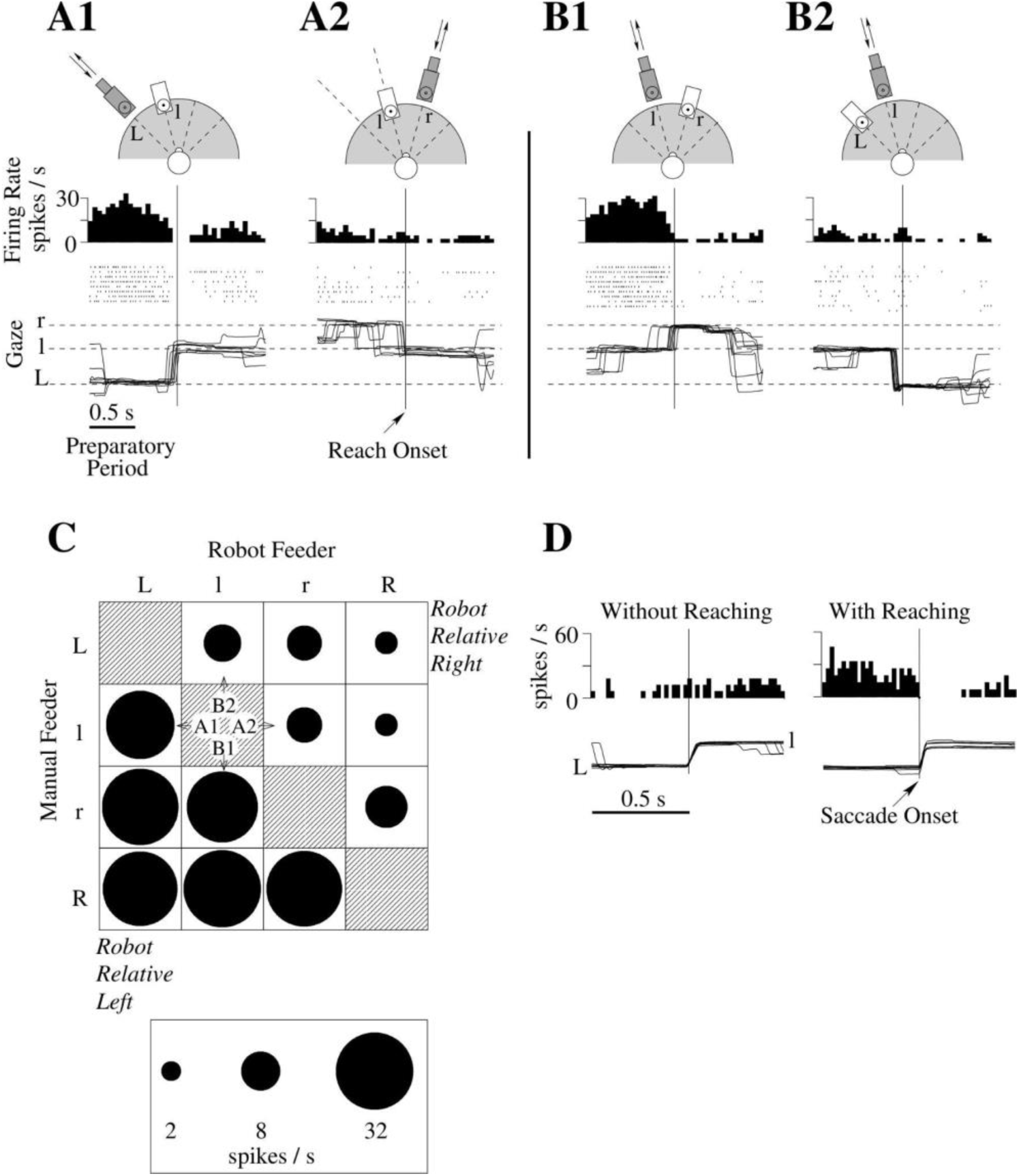
A PFdl neuron with relative-location tuning. **A1**, **A2**, **B1** and **B2** show recordings of neuronal activity and horizontal gaze direction aligned in time on reach onset. Perievent time histograms (PETHs) have a bin width of 50 ms; raster displays consist of lines of dots, with each dot representing the time of a neuronal discharge. Drawings above each display show feeder configuration in the format of Fig. 1. **C**. Firing rates of the same neuron during the preparatory period are illustrated for different feeder configurations using a matrix format. The area of filled circles is proportional to firing rate (see scale below). Prevalence of activity for matrix elements below the major diagonal indicates tuning to the robot feeder being situated to the left of the manual feeder (see Fig. 1). **D**. PETHs centered on rightward saccade onset are shown together with horizontal eye-position records. High firing rates were observed only if the saccade occurred in association with reach (right) and not if it occurred more than 1 s before reach onset (left). Therefore, the tuning properties of this neuron cannot be explained by oculomotor activity alone.

Three task periods were analyzed: (1) an *initial period* preceding the onset of robot movements (0.0–2.4 s from the start of the trial), (2) a *preparatory period* preceding reach onset (0.5–1.0 s before liftoff of the monkey’s hand from the touch pad), and (3) a *reach period* (from 0.1 s before until 0.3 s after liftoff). During the initial period, the monkey alternatively fixated the R and M, during the preparatory period it typically fixated the R and during the reach period it fixated the target of reach, *i.e.,* the R in Task 1 and the M in Task 2. Because of this fixation pattern, eight task periods were analyzed for each neuron, four for each task. Figure 3 shows, for each task, how the initial period was divided into two subsidiary periods depending on the monkey’s fixation point. As summed up in Table 1, these task periods provided variability in gaze angle (to the R or M), orientation of selective attention (to the R until the reach period in Task 2) and reach target location (which varied according to the rule) needed for discrimination among neuronal activity patterns predominantly representing any one of these variables or various combinations.

**Figure 3.**
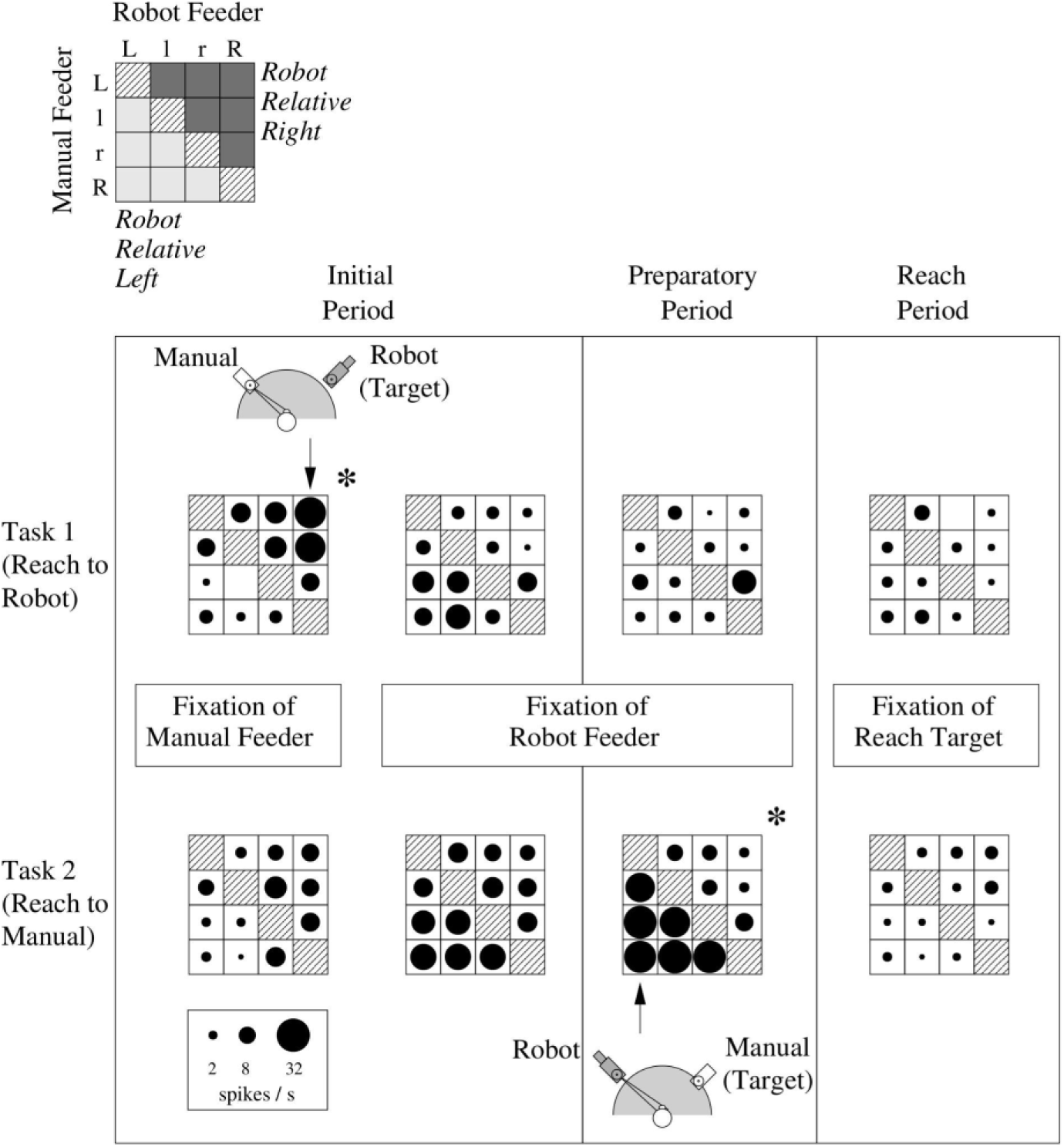
Activity of the same PFdl neuron as in Fig. 2, for both tasks and the four task periods, in the format of Fig. 2C. Matrix columns correspond to the robot feeder’s position and rows correspond to the manual feeder’s position. Task period and fixation conditions are indicated for each case. Statistically significant relative-location tuning (asterisks) was found in two conditions, the two cases in which the monkey looked away from the target of reach.

**Table 1.**
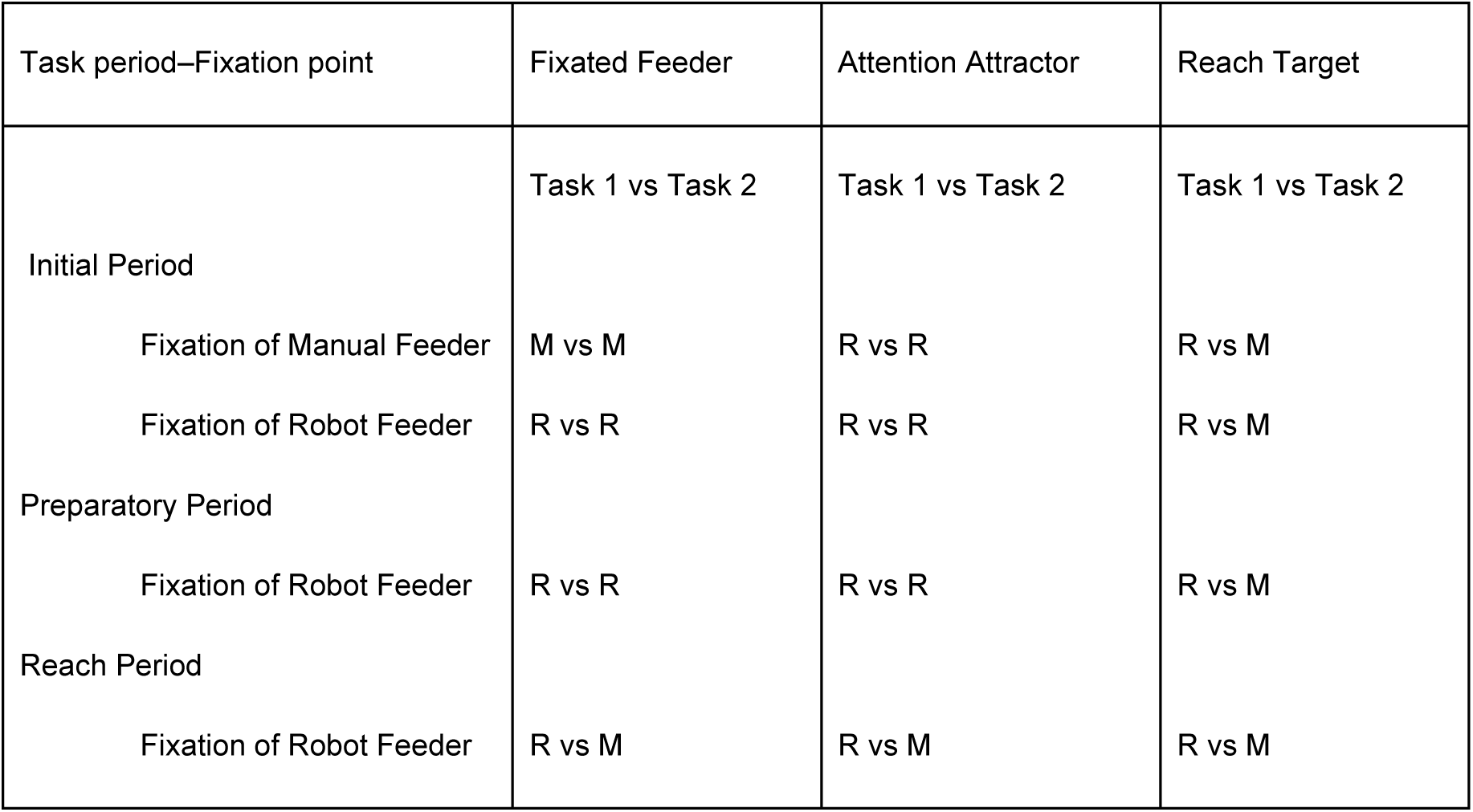
Allocation of eye fixation, orientation of spatial attention and reach target to the manual (M) or robot (R) feeder, depending on task type and period.

Mean firing rates for different configurations of the R and M were represented using firing-rate matrices *F_ij_*, in which rows *i* corresponded to M location and columns *j* correspond to R location (Fig. 1). Note (e.g., Fig. 1 and Fig. 2C) that the major diagonal has no data (hatching) because the R and M never occupied the same location. Each element of the matrix represents the mean firing rate for a given configuration of the two feeders in a given task period. Separate matrices were constructed for each task, task period (initial, preparatory and reach epochs) and fixation condition (toward the R and M during the initial period; e.g., Fig. 3).

A neuron was classified as *relative-location tuned* if and only if it passed three statistical tests. The first test required a statistically significant difference in mean neuronal discharge rate that depended, for any given location of the M, on the R being to the left or to the right of that location. In the first test, therefore, the M served as the *reference* object, the R as the *comparison* object. In the second test, the position of the R served as the reference object. The third test required that the preferences revealed by the first and second tests correspond. Three tests were needed because many neurons were tuned in terms of absolute location of either the R or M, as reported previously (Lebedev and Wise, 2001). For instance, if a neuron reflected absolute location of the R and had a preference for one of the extreme locations (L or R), there would be an average difference between the elements above and below the major diagonal (*F_i>j_* and *F_i<j_*) because the elements for the extreme location would all fall in one of these two categories. However, conducting the three tests resolved this problem: for absolute-location tuning, one of the first two tests would indicate significant tuning, but the other would not. Statistical analysis was performed using a randomization test (Edgington, 1995), which was analogous to ANOVA with factorial design, with relative-location and reference-object position as factors. In this test, statistical significance for the variance was computed using random permutations of trials across the levels of relative location, but within the levels of reference object location.

Once relative-location tuning was detected for a neuron, we classified this property in terms of the spatial variable (target of reach, orientation of spatial attention, or gaze angle) that predominated its relative-location tuning. Eight task-periods were selected for analyses (Table 1, Fig. 3) because they provided the variability of gaze angle, reach-target location and orientation of selective spatial attention needed to discriminate between the tuning patterns related to these variables. We stress, however, that many neurons had mixed properties, and this analysis is only meant to convey a general idea about what most affected the neuron’s relative-location tuning: fixation of the relatively left/right object; attention to the relatively left/right object; or targeting the relative left/right object for the next reaching movement.

To perform this assessment, we compared the firing-rate matrices for different task periods. For this analysis the matrices were constructed in a specific way, illustrated in Fig. 5B. For one kind of matrix, columns and rows represent the fixated feeder and the other (nonfixated) feeder, respectively (depicted in Fig. 5B1; neuronal data in Fig. 4). In other matrices, the columns represented either the attention attractor (vs. the other feeder, Fig. 5B2) or the reach-target (vs. the other, nontarget feeder, Fig. 5B3). Each such matrix was converted into *z*-scores by subtracting its mean and dividing by standard deviation (Fig. 5A), and the degree of relative location tuning was evaluated as

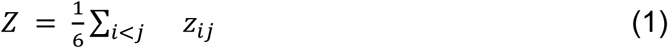

**Figure 4.**
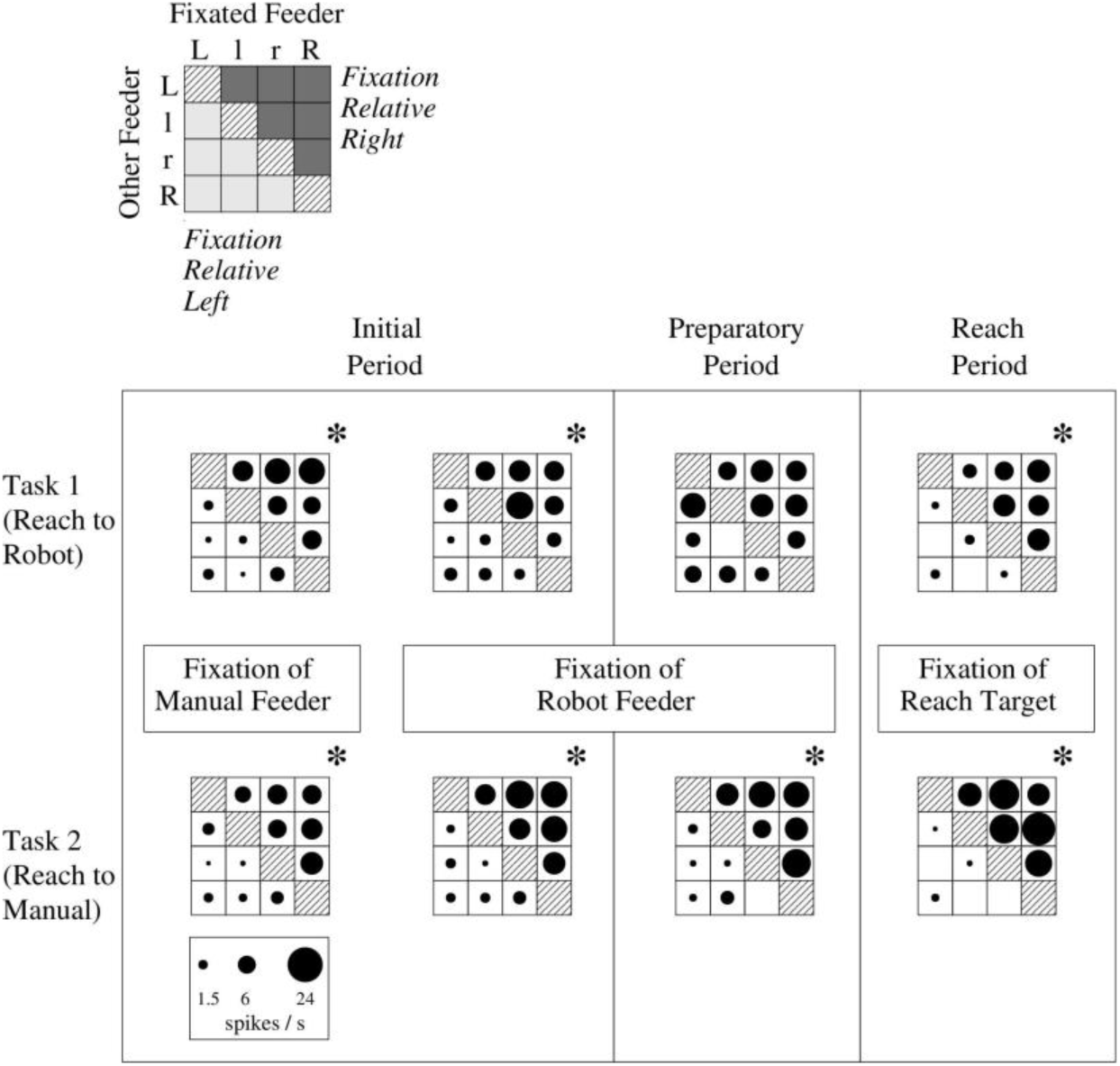
A PFdl neuron with relative-fixation tuning. In each matrix, columns correspond to the location of the feeder at which gaze is directed, whereas rows correspond to the location of the other feeder. Higher firing rates for matrix elements above the major diagonal indicates that this neuron was tuned to fixation of the rightmost of the two feeders, regardless of the feeder identity (R or M). Cases where relative-fixation tuning was statistically significant are marked by asterisks.

**Figure 5.**
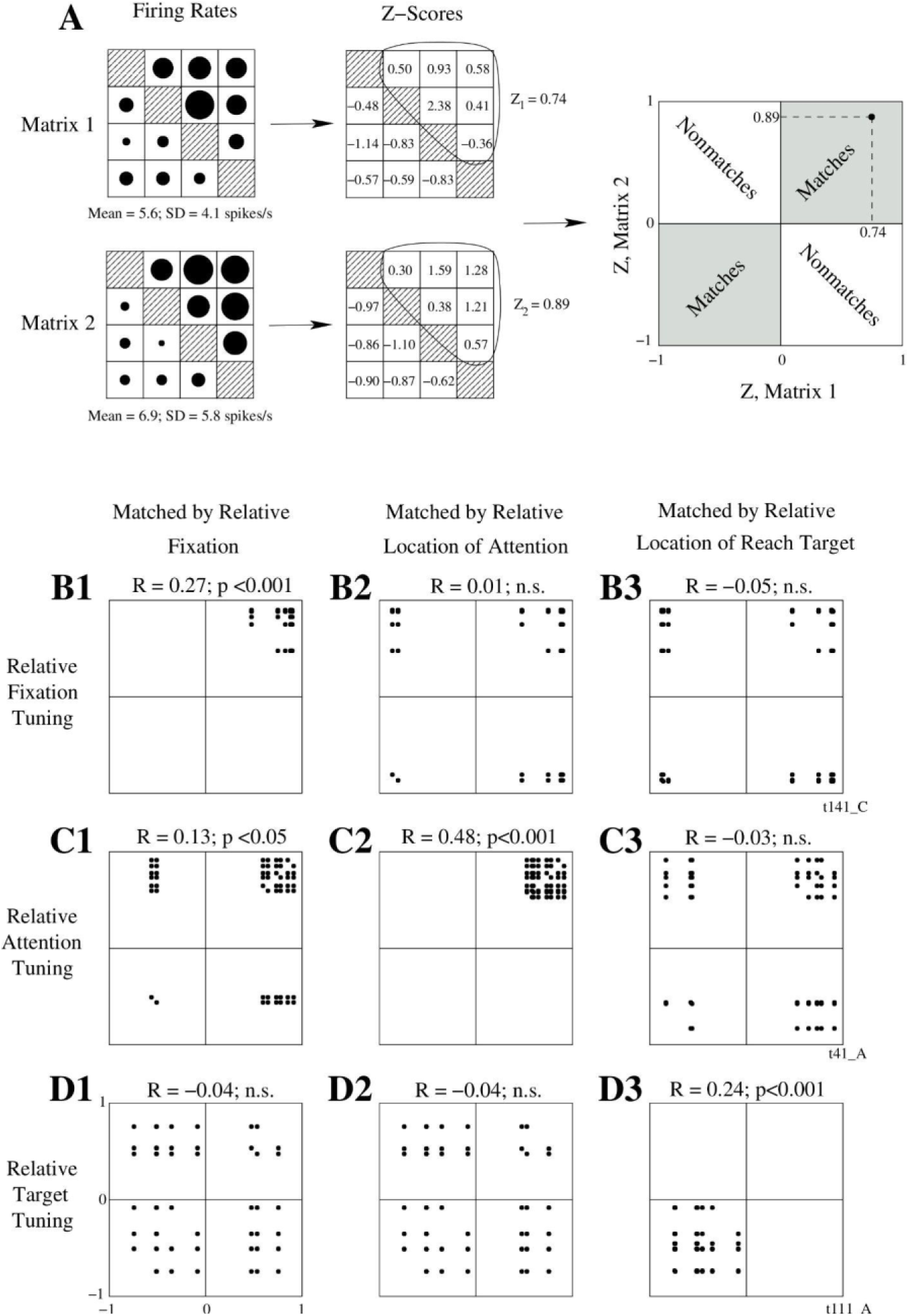
Classification of tuning patterns into relative-location tuning into relative fixation, attention or reach. **A**. Method of calculating *Z*-values as defined by equation 1. Firing rate matrices for two conditions are shown (taken from Fig. 4, initial period, fixation of robot feeder). Each matrix was converted into *z*-scores. *Z* values were calculated as mean *z*-scores for the matrix elements above the major diagonal. High, positive *Z* values for both matrices (0.74 and 0.89) reflect the strength of relative-location tuning. The pair of these *Z* values is represented as a point on an *x-y* plot (top right). Points in the first and third quadrants (shaded) correspond to matching tuning, points in the other two quadrants represent non-matching tuning. In **B1-3**, **C1-3** and **D1-3**, plots were constructed for 3 neurons (B-D) and 3 coordinate systems (1-3): fixation (1), attention (2), and reach (3). *R* values (see equation 2) and their statistical significance are displayed for each of the comparisons. Accumulation of points in the first and third quadrants and high values of *R* correspond to the most plausible classification. One neuron (**B1–3**) had relative-fixation tuning, another (**C1–3**) relative-attention tuning and the third (**D1–3**) had relative-target tuning.

where *z_ij_* are matrix elements expressed as *z-*scores. The absolute value of *Z* corresponds to the degree of relative-location tuning, and its sign indicates directional preference. For example, if rows *i* correspond to the fixated location and columns *j* correspond to the other feeder’s location (Fig. 5B1), then positive values of *Z* indicate directional preference for relative-right R locations (and, of course, relatively left M locations).

To classify neurons, we undertook an analysis of data in these three kinds of firing-rate matrices. This analysis evaluated similarities in relative-location tuning for different task periods and tasks. The degree of matching between the matrices for different conditions was evaluated as

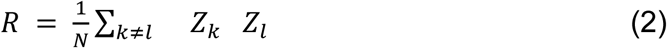

where indices *k* and *l* represent conditions and range from 1 to 8 (*i.e.,* the 8 task periods illustrated in Fig. 3, four task periods for each task) and N is the total number of pairs used to calculate the sum (for 8 conditions used N = 7 x 8 = 56). The requirement *k≠ l* ensures that no condition is matched with itself. If the values of *Z* are high and have the same sign in all 8 conditions (across task periods, fixation points, and tasks), *R* would be positive and high. If they are low and do not correspond in sign, *R* would be close to zero. Statistical significance for *R* was examined using a randomization test in which matrices *z_ij_* were randomly transposed (*i.e.*, indices *i* and *j* were swapped) in equation 1 to calculate statistical distribution for *R*. The analysis of correspondence between the matrices for different conditions is also presented graphically (Fig. 6B-D). In this representation, pairs of *Z*-values (*Z_k_, Z_l_*) are plotted as points. When the matrices matched each other well the points formed a cluster in the first or the third quadrants of the plots; if they did not, clusters of points appeared in several quadrants.

**Figure 6.**
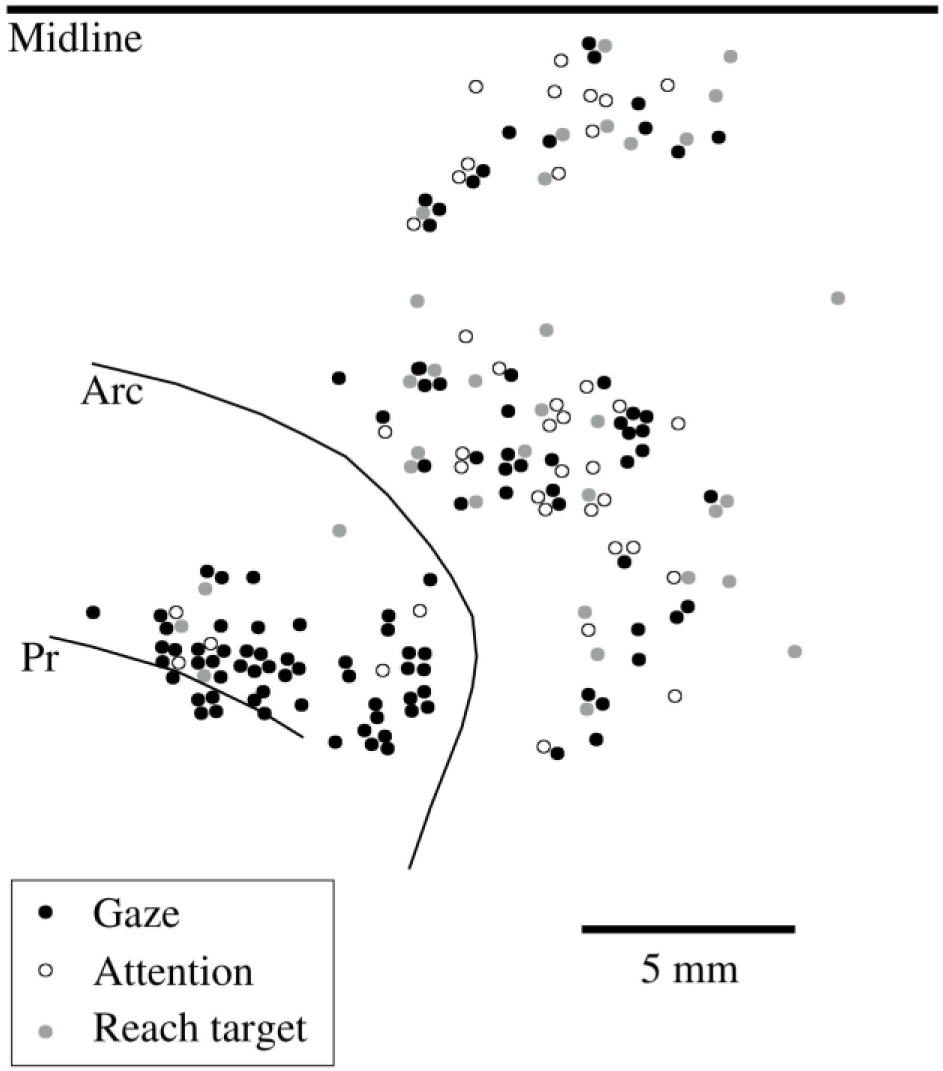
Cortical location of relative-tuned neurons of different types. Information about relative location of gaze, attention and reach target was distributed in the frontal cortex with a prevalence of relative-fixation neurons in PFdl and FEF.

## Results

### Relative-location tuning

Inspection of the firing-rate matrices showed that a substantial proportion of neurons had tuning patterns that depended on the location of the R and M with respect to each other, rather than to their absolute locations in space. These neurons are termed *relative-location-tuned* neurons, in which “tuning” refers to the condition associated with a given neuron’s highest rate of discharge. For example, some neurons were tuned, in this sense, to the R being to the left of the M.

Figure 2C illustrates the firing-rate matrix of such a neuron, one from PFdl. Firing rate data were drawn from the preparatory period in Task 2. Note that the matrix elements below the major diagonal (*F_i>j_*) correspond to configurations in which the R was located to the left of the M, whereas the elements above the diagonal (*F_i<j_*) correspond to relative-right locations of the R (see also Fig. 1). The neuron maintained a high firing rate for all spatial configurations of the feeders in which the R was to the left of the M (*e.g.*, Fig. 2A1 and Fig. 2B1), and its firing rate was less for the configurations in which the R was to the right from the M (*e.g.,* Fig. 2A2 and Fig. 2B2). Accordingly, for neurons tuned to relative-left locations of the R, values of matrix elements below the major diagonal exceeded those for matrix elements above the diagonal. (Note that the reverse holds if one thinks about relative coordinates based on the M, rather than the R, as we do arbitrarily in the above description.)

Figure 2A, B also shows records of horizontal eye position (gaze orientation). The cessation of neuronal discharge occurred shortly prior to the onset of reaching movements, but the monkey made saccades that preceded reaching, as well. During the preparatory period, the monkey typically fixated the R, which by task design served as *attention attractor*, and made a saccade to the M, *i.e.,* the target of reach in this task, Task 2, ∼70 ms before lifting its hand from the touch pad. Because of that temporal correlation, modulations of neuronal activity could have represented oculomotor signals. However, two observations argue against a purely oculomotor nature of the modulations. First, modulations associated with saccades made more than 1 s before reaching had a different pattern (Fig. 2D). Second, this neuron’s activity did not reflect stationary gaze angle. Indeed, starting gaze angles in Figs. 2A1 were equal to ending angles in Figs. 2B2. Similarly, the ending gaze target for Figs. 2A1 was close to the starting one for Figs. 2B2. In both cases, however, firing rates depended on factors other than gaze. Although this neuron did not encode saccades or gaze angle, *per se*, it will be shown below that fixation did have an important role in its activity.

Neurons typically exhibited relative-location tuning during some, but not all task periods. Of the task periods with relative-location tuning, approximately as many were characterized by a preference for the R being to the relative left as to the relative right (306 vs. 283, summed across areas). Table 2 gives the numbers of relative-location tuned neurons in each of the five frontal cortex areas explored. The highest percentage of these neurons was found in PFdl, FEF and PMd (77, 63 and 57%, respectively), and the percentage of relative-location tuned neurons in PMv and SMA was lower (39 and 34%), a difference that was significantly different (χ^2^ = 34.6, n = 1, P<0.001).

**Table 2.** Number of relative-location tuned neurons by task, trial epoch and cortical area. The bottom row shows the percentage of neurons with relative-location tuning in at least one case. Abbreviations: PMd, dorsal premotor cortex; PMv, dorsal premotor cortex; SMA, supplementary motor cortex proper; PFdl, caudal part of the dorsolateral prefrontal cortex; FEF, frontal eye field.

|  | PFdl |  | FEF |  | PMd |  | PMv |  | SMA |  |
| --- | --- | --- | --- | --- | --- | --- | --- | --- | --- | --- |
| Task | 1 | 2 | 1 | 2 | 1 | 2 | 1 | 2 | 1 | 2 |
| Initial Period |  |  |  |  |  |  |  |  |  |  |
| Fixation of Manual Feeder | 29 | 22 | 8 | 5 | 44 | 39 | 8 | 6 | 15 | 7 |
| Fixation of Robot Feeder | 30 | 29 | 7 | 6 | 33 | 26 | 1 | 6 | 5 | 5 |
| Preparatory Period |  |  |  |  |  |  |  |  |  |  |
| Fixation of Robot Feeder | 7 | 23 | 3 | 3 | 15 | 38 | 2 | 2 | 7 | 11 |
| Reach Period |  |  |  |  |  |  |  |  |  |  |
| Fixation of Target Feeder | 22 | 31 | 2 | 4 | 27 | 36 | 6 | 7 | 1 | 9 |
| Relative-Location Tuned/Total | 64/83 (77%) |  | 12/19 (63%) |  | 113/199 (57%) |  | 26/67 (39%) |  | 46/135 (34%) |  |

### Task effects

Figure 3 shows more data from the neuron on which Fig. 2 was based. Note that, during the preparatory period, relative-location tuning occurred only during Task 2, and virtually no tuning occurred during the same period of Task 1. An absence of relative-location tuning during the preparatory period of Task 1 was common. Table 2 shows the percentage of relative-location tuned neurons for different task periods. The number of relative-location tuned neurons was lower during the preparatory period of Task 1 compared to that during the same period of Task 2 (34 vs. 77 neurons; c^2^ = 17.8, n = 1, P<0.001). This was especially clear for PFdl (7 vs. 23; c^2^ = 9.15, n = 1, P<0.002) and PMd (15 vs. 38; c^2^ = 10.6, n = 1, P<0.001), but was less clear elsewhere, in part due to the smaller sample size. Task effects were also significant during the reach period, with fewer neurons showing relative-location tuning during Task 1 than Task 2 (58 vs. 87 neurons; χ^2^ = 6.3, n = 1, P<0.02).

However, this difference was not statistically significant for individual areas. As we take up in the discussion, in Task 2 both feeders were relevant to the monkey’s performance, but in Task 1 only the R was relevant to the task. One variety of the task effect was tuning to relative location in Task 2 and tuning to absolute location of one of the feeders in Task 1. Another involved relative-location tuning in Task 2, but no clear tuning pattern in Task 1. The neuron illustrated in Fig. 3 shows the latter task effect during the preparatory period, and we return below to additional features of its tuning.

During the initial period, the proportion of relative-location tuned neurons depended on fixation conditions. When the monkey fixated the M, more neurons had relative-location tuning in Task 1 than in Task 2 (104 vs. 79; c^2^ = 3.9, n = 1, P<0.05), and when it fixated the R there was no difference (76 vs. 72; c^2^ = 0.07, n = 1, n.s.). To summarize, relative-location tuning was least likely to occur in the preparatory period of Task 1, during which time the monkey was about to initiate reach toward the R, and the M was irrelevant. The greatest relative-location tuning in Task 1 occurred when the monkey fixated the M, even though it was irrelevant. On the other hand, more neurons had relative-location tuning during the preparatory period of Task 2, when the monkey needed to attend most assiduously to the R as it was soon to deliver the go signal by coming within reach, although the reach target was the M. An example of a neuron that followed this schema is shown in Fig. 3 (taken up in detail below).

### Types of relative-location tuning

Relative-location tuning could be related to different spatial variables involved in preparation and execution of reaching movements. We tested whether relative-location tuned neuronns better represented the relative location of (1) the fixation point, (2) the orientation of spatial attention or (3) the reach target. Analyzing activity during a trial single epoch could not distinguish these possibilities, but comparison across epochs could.

Figure 3 illustrates this through further analysis of spatial tuning in the neuron illustrated in Fig. 2. Statistically significant relative-location tuning was detected only for two cases (noted by asterisks above the matrix): the preparatory period of Task 2 (also illustrated in Fig. 2C) and the epoch of the initial period of Task 1, when the monkey fixated the M. During the initial period of Task 1, the neuron was tuned to the relative-right location of the R; during the preparatory period of Task 2, the neuron was tuned to the relative-left location of the R. This pattern of results appears to be complex when viewed from the perspective of the R’s location. However, the monkey fixated different objects in these two cases, and when expressed in terms of relative location of the fixated object, this neuron’s directional preference can be explained parsimoniously: the neuron had a higher activity when the monkey fixated the object that was *not* the target of reach and the target of reach was to the right of the fixation point. These data are not consistent with the neuron encoding fixation of the *leftmost* feeder (it occurs only 2 of the 8 task periods, but the monkey fixates the leftmost feeder for some elements in all of those matrices).

To classify tuning types according to target, fixation or attention relative tuning, we compared the firing-rate matrices following the scheme of Table 1. Figure 4 shows the fixation-tuning comparison for a PFdl neuron. Matrix columns represent the fixated feeder and rows represent the other feeder. All matrices in Fig. 4 show very similar tuning patterns indicating relative-location tuning to the fixated feeder. In all instances, the neuron’s activity for fixation of the rightmost feeder (above the major diagonal) exceeds that for the leftmost feeder (below the major diagonal). Thus, unlike the neuron illustrated in Fig. 3, the neuron shown in Fig. 4 has activity that can be parsimoniously accounted for in terms of the relative location of the fixated feeder, regardless of the fixated feeder identity (R or M).

Figure 5 illustrates the classification of relative-location tuning into relative fixation, attention or reach. In this analysis, the coordinate frame was sought that showed the consistent type of tuning across the 8 task periods. For each type of tuning, values of *Z* were calculated (Fig. 5A), then *R* was computed (see equations 1 and 2). Figure 6B-D shows the plots for pairs of *Z* values for three neurons. Fig. 6B illustrates the same neuron as in Fig. 4. It can be seen that when the matrices were matched by relative fixation, a single cluster of points appeared in the first quadrant of the plot (Fig. 6B1). The value of R was high (0.64) and statistically significant. For the other comparisons, clusters of points appeared in several quadrants, and the values of R were low and not significant statistically. Figure 6C,D illustrate neurons with relative-location tuning for attention and reach target, respectively. For each of them, clusters of points in the first (upper right) or third (lower left) quadrants can be seen in the respective plot (Fig 6C2 and 6D3).

Using this analysis, relative-location tuned neurons were classed into (1) relative-fixation neurons, (2) relative-attention neurons, (3) relative-target neurons, or (4) unclassified neuronls. For a neuron to fall into one of the first three classes, it was required that it had the highest value of R for the respective kind of matrix and that this value was statistically significant (P < 0.05). Otherwise the neuron fell in the unclassified group.

Table 3 shows the results of this classification. Neurons reflecting relative location of fixation targets were predominant in PFdl and FEF (71 and 75% of relative-location tuned neurons in these areas, respectively) and less frequent in PMd (26%), PMv (31%) and SMA (24%) (c^2^ = 16.5, n = 1, P<0.001). Neurons representing orientation of spatial attention in relative terms were found most frequently in PMd (19%), PMv (12%) and SMA (22%) and less frequently in PFdl (6%) and FEF (8%) (χ^2^ = 5.0, n = 1, P<0.03). And neurons representing relative-location tuning to reach target were most frequent in PMd (15%), PMv (15%) and SMA (17%) and less frequent in PFdl (6%) and FEF (0%) (χ^2^ = 4.4, n = 1, P<0.04).

**Table 3.**
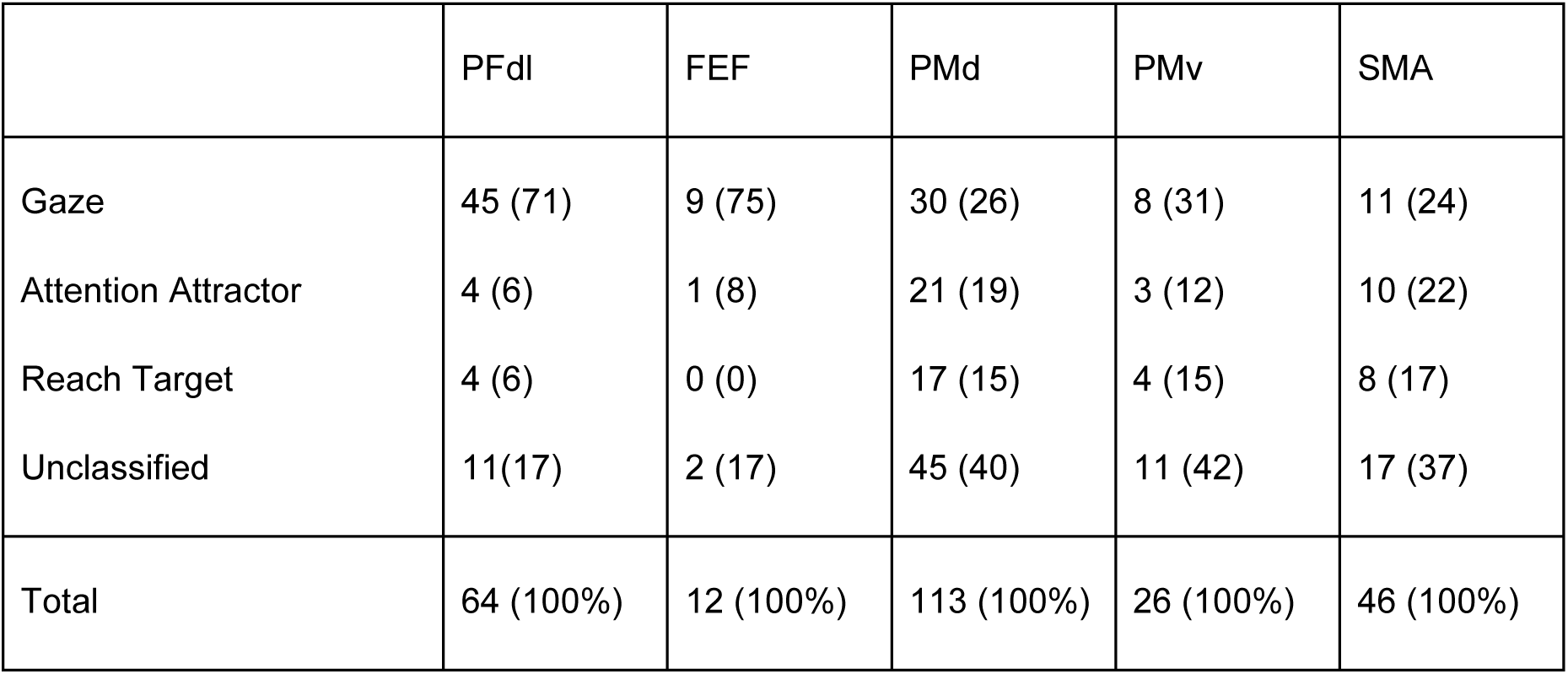
Relative-location tuning types by cortical area. Abbreviations: PMd, dorsal premotor cortex; PMv, dorsal premotor cortex; SMA, supplementary motor cortex proper; PFdl, caudal part of the dorsolateral prefrontal cortex; FEF, frontal eye field.

Note that many neurons fell in the unclassified group, that is they could not be described as simply representing gaze angle, orientation of selective spatial attention or target of reach. The smallest number of unclassified neurons were in PFdl and FEF (17 and 17%), and the proportion of such neurons was high in PMd, PMv and SMA (40, 42 and 37%) (χ^2^ = 11.2, n = 1, P<0.001).

### Cortical distribution of relative-location tuned neurons

We found relative-location tuned neurons throughout the sampled cortical area (Fig. 6). Fixation, attention and reach-target neurons were intermixed with the exception of PFdl and FEF, where relative-fixation neurons prevailed. The other areas contained neurons of all types.

## Discussion

The present study shows that relative-location tuned neurons are plentiful in the frontal cortex of a macaque monkey. Of the areas sampled, relative-location tuned neurons were somewhat more common in PFdl, FEF and PMd (∼66%) than in PMv and SMA (∼37%). Moreover, in the majority of PFdl and FEF relative-location neurons (∼73%), this tuning reflected relative location of fixation, whereas in PMd, PMv and SMA relative-fixation neurons were less numerous (∼27%), and relative-location neurons there reflected the location of attention attractor (∼18%), reach target (∼16%) or more complex tuning patterns. It seems likely based on the present results that the original report of relative-location tuning by Niki (1974) in PFdl probably resulted from the monkey’s fixating the relatively left or right target rather than their status as relative target, as originally thought. Of course, the absence of an oculomotor record in that study precludes firm conclusions along those lines.

We emphasize that most of the examined neurons had complex properties that our classifications, in terms of fixation, attention and reach targets capture only approximately. For instance, the prominence of relative-fixation tuning in PFdl does not exclude participation of the same neurons in the mechanisms of spatial attention or in motor preparation and execution. It is noteworthy that many PMd neurons had complex tuning properties and could not be assigned to any of the simple classes. The comparison between cortical areas is still informative, and it showed that neurons in PFdl and FEF mostly reflected which of the two feeders the monkey fixated, whereas tuning of PMd, PMv and SMA neurons was not as strongly dependent on gaze and reflected the orientation of spatial attention or the target of reaching movement.

Our classification scheme comes with an additional caveat. Suppose that a neuron exhibits a high firing rate each time the monkey fixates the leftmost of the two feeders. According to our classification, this is a relative-fixation neuron with leftward preference. However, this does not necessarily mean that the neuron represents the properties of the fixated object and not the presence of at the periphery. Likewise, suppose that a neuron has increased firing rate when the reach target is the leftmost of the two feeders. Again, this could indicate representation of some properties of the reach target or, alternatively, labeling of the second object as non-target. A neuron representing an object to avoid could be involved in the process of suppression of unwanted, perhaps prepotent, responses. These possibilities need to be investigated with different experimental designs.

### Why relative tuning?

For the monkey to be able to reach toward the target, motor commands must be transformed eventually into patterns of muscle activation, for which the relative locations of objects has little, if any, relevance. A possible role for relative-location tuning is, then, involvement in oculomotor control and/or perception and/or decision making. The relative location of the two highly salient objects, the feeders, can be parsimoniously encoded by reference to each other, and this approach might have computational advantages over an absolute reference frame. Indeed, if only two locations matter for a given behavior, representing space in relative terms could aid efficient decision making. Thus, a response choice could be formulated by a higher-order brain area as “reach toward the rightmost object”. Studies on cortical representation of both eye movements (Olson and Gettner, 1995; Olson and Tremblay, 2000; Tremblay et al., 2002) and food value (Tremblay and Schultz, 1999; Hikosaka and Watanabe, 2000) support the idea that frontal cortex plays a role in relational coding for decision making. Indeed, reference frames based on absolute coordinates (*e.g.,* retinocentric maps) and those based on the relative location of gaze, attention, and reach targets could coexist. Many of the neurons in the present neuronal sample exhibited relative-location tuning during some, but not all task periods. Overall, the examined frontal cortex areas appear to integrate the relative and absolute spatial information and do so in a flexible, task-dependent way.

### Behavioral rule and relative location tuning

Task 2 led to stronger relative-location tuning than Task 1, most likely reflecting the fact that in the former both feeders were relevant, whereas in the latter only the R mattered for task performance. However, clear-cut differences were only seen during the preparatory period. During that task period, the disparity between the feeders was most critical because the monkey had to devote considerable attentional resources to the R while preparing a motor response toward the M. Curiously, relative-location tuning strengthened during one period of Task 1: the monkey occasionally reoriented gaze from the behaviorally important feeder, the R, to the irrelevant M, and many neurons exhibited relative-location tuning. Such behavior could be called curiosity or exploration. The enhancement of relative-location tuning during this behavior was particularly strong for SMA, an area thought to be important for self-initiated behaviors (Okano and Tanji, 1987; Mushiake et al., 1991; Halsband et al., 1994; Chen et al., 1995; Thaler et al., 1995).

### Relative-fixation tuning

Relative-fixation tuning was observed in all studied areas of the frontal cortex, but most frequently in PFdl and FEF. Neurons with this property reflected whether the monkey fixated left- or the rightmost of the two feeders. An straightforward explanation of this type of tuning would be that it was simply related to eye movements. However, the neurons tuned this way did not reflect the angle of the eye in the orbit because for a given eye position they could have high or low rate, depending on the feeder arrangement (Fig. 2A, B). Relationship to the prepared saccade direction was not a universal property, either (Fig. 2D). Additionally, the relative-fixation tuning depended on the behavioral task and task period.

Although not directly comparable, our properties of relative-fixation neurons in the frontal cortex resemble the properties of gaze dependent neurons in PPC (Batista et al., 1999). The PPC neurons were active prior to the reaching movements, and their activity depended not so much on the initial hand location as on initial gaze orientation. Batista and his colleagues interpreted these neurons as encoding reach direction in retinocentric coordinates. Since frontal and parietal cortices are richly interconnected (Jones and Powell, 1973; Johnson et al., 1996; Marconi et al., 2001), it would not be surprising to find neurons with similar properties in the frontal cortex.

The computational theories of reaching employ the concept of a difference vector, that is the difference between the current locations of the limb and movement target (Bullock and Grossberg, 1988; Guenther et al., 1994; Cisek et al., 1998; Burnod et al., 1999). Neurophysiological evidence supports this idea (Buneo et al., 2002), as well as the modeling (Deneve et al., 2001; Andersen and Buneo, 2002). This theory holds that reaching movements are planned in terms of the relative location of the target of reach and the hand’s current location. Also according to this theory, this difference vector arises among PPC networks by subtraction of two vectors, one from some origin to the target, the other from the same origin to the hand’s location. This computation is thought to use retinocentric coordinates and is assumed to have as its origin the fovea. When viewed from this perspective, the neuron illustrated in Fig. 3 has activity that is consistent with representing the first of these two vectors, the vector from the origin to the target. When the monkey fixates the target, the neuron has relatively little activity. However, during the task period involving fixating the nontarget feeder, the neuron shows a clear preference for the target locations to the right of the fixation point (marked with asterisks in Fig. 3). Furthermore, when the monkey fixates the R during the initial period of Task 2 and looks at the R but before it begins moving, there is high activity both above and below the major diagonal. This could occur because the reach target is to the right of gaze (for data below the major diagonal) and the attention attractor is to the right of gaze (for data above the major diagonal). Although speculative, this account of activity such as that illustrated in Fig. 3 sums up the properties of this complex neuron in a reasonably parsimonious manner.

However, not all of relative-fixation neurons had this pattern of activity. Many showed relative-fixation tuning irrespective of whether the monkey looked at or away from the target of reach (Fig. 4). Thus, the neurons that could have contributed to encoding the target in retinocentric coordinates constituted only a part of the present sample.

We found relative-fixation tuning in a substantial proportion (∼27%) of premotor neurons. The influence of gaze orientation on movement-related activity has been reported in PMv (Boussaoud et al., 1993; Mushiake et al., 1997), PMd (Boussaoud et al., 1993; Boussaoud, 1998; Boussaoud and Bremmer, 1999; Cisek and Kalaska, 2002), SMA (Fujii et al., 2002), and the pre-supplementary motor area (Fujii et al., 2002). Neuroimaging results in humans also suggest that mismatches in gaze and reach targets generate higher neural signals in both frontal and posterior parietal cortex (Baker et al., 1999; DeSouza et al., 2000). Some of these effects could be attributed to relative-fixation tuning. This idea helps resolve an apparent discrepancy between the report that emphasize gaze dependency and those that minimize it (Fogassi et al., 1992; 1996; Graziano and Gross, 1998; Cisek and Kalaska, 2002). Many of the latter studies have been based on receptive-field-like analyses of neuronal activity, in which minor differences in activity receive less attention than the overall region in which stimuli drive the neuron above background levels of discharge. The modest, quantitative effects of gaze that have been reported for the frontal motor areas is what would be predicted if the motor plan results for a difference-vector computation in retinocentric coordinates.

### Relative-attention tuning

To class neurons as reflecting the relative orientation of spatial attention, we required that they do so regardless of fixation. This neuronal population is probably much smaller than that actually involved in the orientation of spatial attention. However, even with our conservative criterion, ∼18% of premotor cortex neurons with relative-location tuning were classed as involved in gaze-independent spatial attention. We previously reported that PMd neurons represent orientation of spatial attention (Lebedev and Wise, 2001), and the present finding extends this observation to relative-location coding.

### Relative-target tuning

Neurons representing the relative location of reach target were found predominantly in premotor areas: PMd, PMv, and SMA. These areas are thought to be crucially involved in movement planning. However, even in these areas the proportion of such neurons among all those reflecting relative-location tuning was somewhat low (∼15%). This finding could result, in part, from the stringent requirement that a neuron represent reach target throughout all task periods, including the ones preceding the movement by several seconds. It is possible that some neurons began to represent the target only immediately before the movement, and such neurons escaped classification according to the present analytical method. However, to distinguish between neurons reflecting the relative location of a reach-target neurons and those reflecting the relative location of spatial attention, we had to use stringent requirements. In addition, we required that during the movement period target neurons were tuned to the same direction as during the preceding periods. These requirements certainly led to an underestimate of the proportion of neurons selective for the relative location of the reach target. Finally, some of the relative-fixation neurons could have contributed to target encoding using a retinocentric reference frame, as discussed above.

### Unclassified tuning

Neurons that had relative location tuning, but could not be classed as representing fixation, target of reach or spatial attention were especially numerous in premotor areas (∼40%). These hybrid neurons were probably attuned to more than one spatial parameter. Such neurons are candidates for representing spatial variables in a distributed neural network.

## Acknowledgements

We thank Mr. Robert Gelhard and Mr. Alex Cummins for preparation of the histological material and Dr. Andrew Mitz for engineering the experimental apparatus.

